# Multiplex staining by sequential immunostaining and antibody removal on routine tissue sections

**DOI:** 10.1101/148742

**Authors:** Maddalena Maria Bolognesi, Marco Manzoni, Carla Rossana Scalia, Stefano Zannella, Francesca Maria Bosisio, Mario Faretta, Giorgio Cattoretti

**Author notes:** equally contributed (Maddalena Maria Bolognesi, Marco Manzoni, Carla Rossana Scalia). Mailing address: Prof. Giorgio Cattoretti, Anatomia Patologica, UNIMIB and Ospedale San Gerardo, Via Pergolesi 33, 20900, Monza (IT).

## Abstract

Multiplexing (mplx), labeling for multiple immunostains the very same cell or tissue section in situ, has raised considerable interest. The methods proposed include the use of labelled primary antibodies, spectral separation of fluorochromes, bleaching of the fluorophores or chromogens, blocking of previous antibody layers, all in various combinations. The major obstacles to the diffusion of this technique are high costs in custom antibodies and instruments, low throughput, scarcity of specialized skills or facilities.

We have validated a method based on common primary and secondary antibodies and diffusely available fluorescent image scanners. It entails rounds of four-color indirect immunofluorescence, image acquisition and removal (stripping) of the antibodies, before another stain is applied. The images are digitally registered and the autofluorescence is subtracted. Removal of antibodies is accomplished by disulphide cleavage and a detergent or by a chaotropic salt treatment, this latter followed by antigen refolding. More than thirty different antibody stains can be applied to one single section from routinely fixed and embedded tissue. This method requires a modest investment in hardware and materials and uses freeware image analysis software. Mplx on routine tissue sections is a high throughput tool for in situ characterization of neoplastic, reactive, inflammatory and normal cells.

## INTRODUCTION

An increasing percentage of diagnosis in pathology are finalized to the identification of specific proteins, relevant for the patient’s management, via antibodies and the deposition of a pigment or fluorochrome at the protein’s location in the tissue section.The variety of specific antibodies is of increasing numerosity and complexity, ranging in the hundreds for an average lab, targeting constitutive tissue elements, oncogenes, growth factor and hormone receptors, aberrant product of genomic aberrations, etc. Some assays are aimed at identifying diseases or refine the pathologic classification, others at guiding the therapy. The standard is to perform one stain at the time, on serial sections, until the amount of material allows it and no competitive tests (e.g genetic or extractive) are favoured over the immunostain (1). In the life sciences realm, fluorescent reporters are used more often, not necessarily viewed under a microscope, but also with a variety of other quantitative methods (flow cytometry, image cytometry, confocal microscopy etc.) which better suit the analysis of live or lightly fixed cells. The fundamental difference between immunohistochemistry (IHC) and immunofluorescence (IF) is the use of light: absorption for IHC, emission for IF. It is connatural with IF the diversification of fluorescent reporters, limited by the source of light, the overlap of fluorescence spectra, the available fluorochromes and, ultimately, the cost of the reagents.

Surgical Pathologists do not venture outside IHC and if so, very rarely (2). Life science scientist follow the ground rules of IHC when dealing with fixed and embedded tissues (3).

Recently, the interest in performing multiple assays on formalin fixed, paraffin embedded (FFPE) specimens has gained ground, both in Pathology and in the Life Sciences field and is currently referred to as multiplexing (mplx). In order to set aside a plethora of multiple staining methods of this type of material already published (4-6), we consider only methods in which more than three different stains are performed on the very same slide.

A strategy to perform multiple stainings at the same time on the same section involves the use of animal-specific secondary antibodies linked with a reporter directed against antibodies raised in different animals (rabbit, goat/sheep, rat and three-four mouse isotypes). Given the limited variety of antibody sources, it is inevitable to devise mplx strategies to stain antibodies raised in the same species with contrasting colours. This has been accomplished essentially in three ways: *i)* bleaching directly conjugated primary antibodies before adding other layers, *ii)* blocking the access to a previously deposited antibody for a second staining round, or *iii)* removing antibodies from sections after staining and imaging.

The use of directly labelled primary antibodies has been pioneered by Schubert et al. (4), followed by others (7-9). After the tissue image has been acquired, the fluorophore is inactivated by UV light (4), alkaline solutions (8) or sodium borohydride (7). The major drawbacks of this technology is the cost of the directly labelled antibodies, the requirement for a customized conjugation, the loss of antigens (10) and the occasional lack of sensitivity for important makers (9, 11). In addition, if a robotized solution is available for otherwise highly repetitive tasks, the number of staining stations is a limiting factor for throughput (4, 9). Lastly, the steric hindrance of a previously deposited antibody against the subsequent deposition of the very same antibody has not been addressed at all (9, 12).

Blocking the access to the first antibody layer is an old technique (13), based on the blocking ability of an insoluble (and therefore hydrophobic) precipitate such as diaminobenzidine (DAB), which prevents another staining round to get access to the first, even if the antibodies in both rounds are raised in the same species. Since the amount of insoluble precipitate conditions the access of the second staining to the same subcellular or cellular structure, a variety of hybrid IHC-IF methods have been employed (14), using extensive antibody titrations and differential reporter sensitivity, e.g with tyramide (insoluble) precipitation (15).

A purely immunologic block by using Fab monomeric fragment dates back several years (16, 17) and has been used rarely for multiplexing in IF: concentrations of the Fab fragments in excess of 500 μg/ml, required to achieve a non-complete blocking, are costly and impractical (17, 18).

Removal of the previously deposited layer of antibodies is a strategy tested by multiple investigators (8, 14, 19-23) and involves a broad range of solutions, chemical agents and temperature. The protocols range from a very mild boiling in Antigen Retrieval solution (15) (shown by us (20) and others (24) to be inefficient at removal of antibodies), to an acidic Glycine buffer (23), to strong chemicals (22), to proteases (8) and to a mixture of strong reducing agent and a detergent (19, 20).

The cyclic deposition of an immune layer, the capture of the image, the removal of the antibodies and reporters and a new immunostaining is suitable both for IHC and IF methods (14, 19, 20).

Alcohol-soluble precipitates and single color IHC are required for sequential IHC staining. In fluorescence, the limits are dictated by the fluorochromes and the filters. Additional colors may be added to the standard four (DAPI, FITC, TRITC, Cy5) by using spectral deconvolution of the fluorescent signals (25, 26).

Most of these methods for multiplexing had a limited diffusion and only a few could demonstrate more than a dozen staining on the same section, with the exception of two methods based on directly labelled antibodies (4, 9), only one applied on routinely treated sections.

The landscape of mplx may change because of the introduction of the so-called “next generation IHC”, a multiplex technique based on isotope-tagged antibodies and in situ mass-spectrometry detection (reviewed in (27)). In addition, barcode labelled probes, including antibodies, may allow quite extensive multiplexing with the NanoString technology (<u>http://NanoString.com</u>). Though, the ability to visualize single cells in a whole slide image of both systems is unknown.

Having published a preliminary evidence for a potentially high volume multiplexing by antibody removal (20) and established the foundation for the reproducibility of the method (28), we embarked in a study to investigate antibody removal methods, one novel, and multiplexing for >30 antibodies on a single routinely processed section.

## MATERIALS AND METHODS

### Tissues and Antigen Retrieval

Formalin fixed, paraffin embedded (FFPE) fully anonymous human leftover material used was exempt from the San Gerardo Institutional Review Board (IRB) approval as per Hospital regulations (ASG-DA-050 Donazione di materiale biologico a scopo di ricerca e/o sperimentazione, May 2012).

Three micron sections were cut and placed on coated glass slides (SuperFrost^R^Plus, Bio-Optica, Milano, IT), baked in a vertical position overnight at 40C or for 1 hour at 60C, dewaxed using xylene, rinsed in a graded alcohol series, rehydrated in distilled water (29); antigen retrieval (AR) was performed as published (30) with (ARx) or without (AR) as pre-treatment. Sections in distilled water were inserted into radiotransparent slide holders (model #S2029; Dako, Glostrup, DK) and transferred to a glass container filled with 800 mL of the retrieval solution (10 mM EDTA in Tris-buffer pH 8) (Sigma-Aldrich, Milan, Italy). The container was irradiated in a household microwave oven at full power for 8 minutes, followed by 20 minutes of intermittent electromagnetic radiation to maintain constant boiling. Sections were cooled to about 50C before transferring to buffer, being ~60C a sort of landmark between two temperature ranges with quite different effect on antigens (30). The ARx procedure maximize the antigen exposure in tissue, overcoming the requirement for a pH-dependent retrieval for individual antibodies (30).

Sections not being stained for extended periods of time (typically > 3 days) were stored at -20C in 50% glycerol (Sigma-Aldrich) in pH 7.5 Tris buffer and 300 mM sucrose (5% of a saturated solution).

### Primary and secondary antibody dilution and incubation

Primary antibodies to be used for multiplexing (Supplemental Table 1) were screened for sensitivity and specificity (31) on target positive tissue sections at 1 and 0.1 μg/ml dilution in TrisHCl Buffered Saline (TBS) to which 2% BSA, 0.05% Sodium Azide and 100 mM Trehalose were added, counterstained with an Alkaline-Phosphatase conjugated secondary antibody and developed with nitro-blue tetrazolium and 5-bromo-4-chloro-3′-indolyl phosphate (NBT-BCIP, Roche) (32). If the antibody concentration was not known, two one-log dilutions of the recommended dilution were tested. One μg/ml was almost invariably the dilution of choice and a saturating concentration. To test for saturating concentration, a 12 ml, 1 μg/ml solution of CD79a in a vertical five slide mailer (model 715409, Electron Microscopy Science, Hatfield, PA) stained 36 sequentially placed tissue sections with an average staining variation intensity of -4% ±6% from time zero.

The dilution and the diluent was found appropriate for both immunofluorescence (IF) and immunohistochemistry (IHC), for either 1 hr or overnight incubation. Compared to 1 hr, overnight incubation increases the staining strength of a factor between 30% and 200%, depending on the antibody (Supplemental Fig. 1-A). Secondary antibodies (Supplemental Table 2) were diluted in the same diluent and used at the 1:300 dilution.

Incubation was at room temperature in a horizontal plastic slide box (Kartell, Italy) containing a moist paper towel. At least 100μl of antibody was applied for a section of 1x1 cm or less and the volume multiplied accordingly for larger sections.

TBS buffer to which 0.01% Tween-20 (Sigma) and 100 mM sucrose were added (TBS-Ts) was used throughout all the experiments for washing. Three 5 min changes of buffer were used throughout.

For IF, antibodies of different isotype and/or species were pooled, each at the final dilution and incubated for the designed amount of time.

After washing, pooled, non-crossreactive conjugated secondary antibodies were applied for 30 min.

A shortened sequence of primary and secondary antibodies (30 min each, with three TBS-Ts washes in between), so called double indirect IF staining, repeated once after the first IF staining cycle (32) doubles the staining intensity (Supplemental Fig. 1-B) and was used when appropriate.

The variability of the staining efficiency, measured on duplicate parallel staining in serial sections, averages 3.1% (min 0%, max 12.3%) of a given fluorescent channel and is shown in Suppl. Table 3.

Slides stained in IF were mounted with Phosphate buffered (pH 7.5) 60% Glycerol -40% distilled water mixture containing 0.2% N-propyl Gallate and 584 mM sucrose (10% of a saturated solution), to which DAPI dilactate 5.45 μM (Sigma) were added, the latter from a 1.09 mM stock solution in PBS. The concentration of DAPI was adjusted so that the DNA DAPI fluorescence did not bleed into the other channels.

Preliminary work showed that a Glycerol-based mounting medium containing sucrose maintain or increases the antigen availability, differently from a Polyvinyl alcohol mounting medium (Sigma) containing 10% sucrose or other mounting media (e.g. Gycerol-Gelatin, C0563, Dako; FluoroGel, Electron Microscopy Sciences, Hatfield, PA), which decrease antigenicity (Supplemental. Fig. 1-C). Hardening fluorescent mounting media and glycerol-gelatin mix were found to cause antigen re-masking with repeated use (not shown).

### Antibody stripping

Coverslip was gently removed by soaking the slides in TBS or distilled water (no effect on subsequent stripping detected; not shown).

Beta-mercaptoethanol / sodium dodecyl sulphate (2ME/SDS) stripping was performed as published (20), modified by halving the concentration of the buffer with an equal volume of distilled water. Afters stripping, the slides were washed in TBS-Ts for at least 30 min with repeated buffer changes.

Chaotropic salt-dependent removal of antibodies and antigen renaturation was performed as published by Narhi LO et al (33, 34). Sections were immersed for 10 min at 40C in 12 ml 6M Guanidinium Hydrochloride (GnHCl, Sigma) solution in vertical slide mailers. The GnHCl solution was buffered with 0.05M Citric Acid – Sodium Citrate Buffer Solution (0.12 g Citric Acid Monohydrate, 0.17 g Trisodium Citrate Dihydrate in 100 ml).

The 40C temperature was chosen over room temperature for consistency; no other temperatures were tested.

A second shorter passage in 6M GnHCl to which 5% w/v sucrose (290mM) was added, ensured the removal of eluted antibodies. Subsequently, the sections were transferred to 6M Urea solution containing 5% sucrose at 40C for 10 min, followed by another 10 min incubation in 3M Urea and 5% sucrose solution. Finally the sections were left to re-equilibrate in TBS-Ts. In some experiments, sections were then transferred to 10mM EDTA in Tris-buffer pH 8 and irradiated until reaching the boiling for 1 minute and left to cool below 50C.

A schematic depiction of the staining and stripping sequence is shown in Table 1.

**Table 1.**
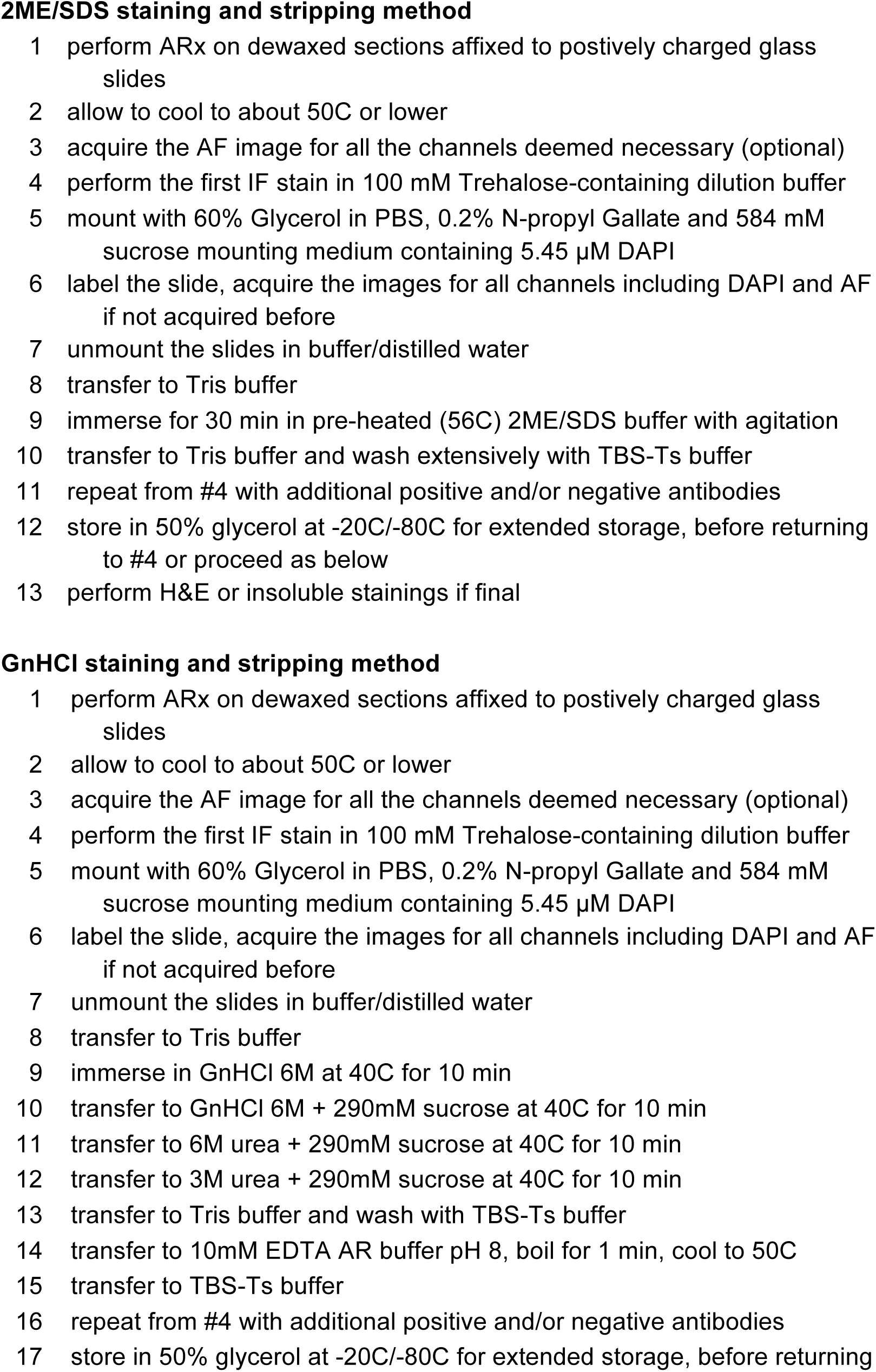
Flow chart of cyclic antibody staining and removal with 2ME/SDS and GnHCl The sequential steps for repeated antibody staining and removal with 2ME/SDS and GnHCl are listed. Abbreviations: 2ME/SDS: 2-mercaptoethanol/ sodium dodecyl sulphate stripping buffer; GnHCl: guanidinium hydrochloride; ARx: two-step antigen retrieval method (30); AF: autofluorescence; IF: immunofluorescence; TBS-Ts: Tris buffered saline containing Tween-20 and sucrose. For detailed composition of buffers and solutions see text.

Removal of the bound immunoglobulins with a sodium borohydride treatment (35) was performed as follows; a stable 4.4M Sodium Borohydride (NaBH_4_) (Sigma) stock solution was obtained by dissolving 1.67 gms in 10 ml of 14M NaOH (Rohm & Haas, <u>https://www.scribd.com/document/326437122/Sodium-Borohydride-Digest</u>).

For antibody removal, sections were equilibrated in 0.05M Tris buffer pH 9, exposed to various freshly made concentrations of NaBH_4_ in the same buffer for 10 minutes, with or without 0.5% SDS, followed by immersion in 0.1M citrate buffer pH 4 to inactivate NaBH_4_.

### Immunofluorescence scanner and virtual whole slide acquisition

The Hamamatsu Nanozoomer S60 scanner (Nikon, Italia) is equipped with an Olympus 20X/0.75 PlanSApo objective, a Fluorescence Imaging Module equipped with a L11600 mercury lamp (Hamamatsu), a linear ORCA-Flash 4.0 digital CMOS camera (Hamamatsu) and two six-positions filter wheels, one for excitation, the other for emission filters, and a three-cube turret. The excitation filters (all from Semrock, Inc, Rochester, NY) are 387/11 (DAPI), 420/10 (AF) 480/17 (FITC), 556/20 (TRITC), 650/13 (Cy5). The emission filters are, in the same order, 435/40, 530/55, 520/28, 617/73 and 694/44. Three dichroic mirrors are: a triband FF403/497/574-Di01 and two single pass Di03-R488 and FF655-Di01. The AF and Cy5 dichroic and emission filters are housed in a cube each. The filter combinations and the fluorochrome excited are accessible at the Semrock Searchlight website; <u>https://goo.gl/dnYoMc.</u>

After the first submission of this manuscript, we became aware that two commercial sources (Chroma Technology Corporation, VT, USA and Semrock, NY, USA) have produced a filter set tailored for BV480 (excitation 436 or 438 nm, emission 478 or 483 nm, dichroic 458 nm) and which may be also used for AF acquisition.

Stained slides to be acquired were labeled with a 2D barcode for automatic file name acquisition (TEC-IT Datenverarbeitung GmbH, Steyr, Austria), the bottom surface rinsed in distilled water to remove traces of sucrose and scanned in batch mode with pre-defined parameters. The barcode content is the sample identifier and the biomarker sequence, each biomarker flanked by a short fluorescence channel acronym for automatic coupling of each image with the biomarker by an image analysis software. The file name was kept as short as possible (i.e. below 20 characters).

Histograms data of fluorescence images as 8bit gray levels (0-255) and of at least 2mm x 2mm were obtained with Fiji and exported in an Excel spreadsheet. Cumulative percentage pixel numbers of the total in each channel was accrued over 255 channels and the channel position where 90% of the pixels are found was used as the measurement of the fluorescence intensity (Scalia et al. (29), supplemental Fig. 2, ibid)

### Autofluorescence

The emission of tissue autofluorescence was quantified by exposing dewaxed, antigen-retrieved reference tissues (placenta, kidney, skin, etc.) to multiple exposure times in each of the filter combinations available (Supplemental Fig. 2-A).

AF was acquired either with a separate channel on four-color stained sections during the image acquisition or on antigen retrieved unstained sections, mounted with DAPI and coverslipped, in each filter combination, before the first round of immunostaining. In this case, the images were acquired at submaximal intensity for AF-rich tissues such as kidney or liver.

The mean pixel AF data for each tissue, each fluorescence channel and each time point was extracted with the Histogram function of ImageJ from the 8bit greyscale images, then plotted as intensity over exposure time in an Excel spreadsheet.

### Chemical inactivation of AF

For endogenous fluorescence quenching, a 1:280 dilution of a sodium borohydride stock solution (NaBH_4_ 15.7 mM, NaOH 0.05 N) was added to either 95% ethanol or to a 0.05 M Tris-buffer pH 9. As a control, an identical dilution of a stock 14M NaOH solution was used. Sections were exposed to the NaBH_4_ or control solution for 10 min and 30 min, either after the 99% ethanol deparaffinization step, or before antigen retrieval. Autofluorescent tissue samples, exposed to NaBH_4_ were imaged with three filter combinations (420/10: AF; 480/17: FITC; 556/20: TRITC) and the autofluorescence values imported in an Excel sheet.

### Digital Subtraction of AF

AF was subtracted essentially as published by Pang et al and Van DeLest (36-38).

AF in some tissues (placenta, lymphoid tissue) had values of the same scale of intensity in each channel where was measured and compared pixel by pixel (Supplemental Fig.2-C and 2-D). In these cases, the linear equations for fluorescence in the AF filter and in the other filter were used to calculate an accomodation factor, in order to equalize the image obtained with the AF filter with the exposure and fluorescent response of the specific image, before subtraction.

As an example, the slope of the regression line for AF excited by the 420 nm filter is 0.61x *x*+2.7=*y*, the slope for the 488 nm excitation is 0.26x*x*+0.31=*y*.

If AF was acquired on placenta at 112ms and the FITC image at 80ms, then the factor to be used would be calculated as: (0.61x112+2.7)/(0.26x80+0.31)=0.29, being the first term the value for AF, the second term the value for FITC and 0.29 the factor to apply to the AF image to equal the AF background in the FITC image.

After importing in Fiji with the BioFormat Plugin all the .ndpi files and saving them as .tiff files, the AF image was adjusted for the numerical factor of accomodation and subtracted from the stained image by using the Image Calculator function.

The procedure worked also in tissues with channel-specific AF values, although manual adjustment of the accomodation factor may be required. Alternatively, pre-staining AF in each specific channel was acquired and then aligned and subtracted.

An example of AF subtraction is shown in Supplemental Fig. 2-B.

AF subtracted images were used throughout for quantitative analysis and for imaging.

### Registration

Removing and replacing the very same slide on the scanner stage entails microscopic translations and rotations, which misalign subsequently acquired images.

To re-align the images (register), one image of the nuclei in the tissue (DAPI) was set as reference; then each other DAPI image acquired with a subsequent scan was aligned with the TurboReg Fiji plugin (39). The coordinates of the registration were recorded as landmark (a .txt file with coordinates) and applied to the entire stacks made up of the images of the same round with the Multi StackReg Plugin (40).

Care was taken to produce whole slide images (WSI) as close to each other as possible; the registration process is considerably more difficult with images of different size, texture and content.

### Image Analysis

The amount of positive pixels, expressed as percentage of the area analyzed, was measured on each grayscale IF staining whole image, after application of an uniform threshold algorithm (Huang), for each staining cycle for that marker. WSI from routine immunohistochemical stain on serial sections, stained with DAB and counterstained with Haematoxylin, were deconvoluted as published (30), inverted and the pixel area quantified with the very same algorithm used for the IF images. Three separate fields per stain were analyzed.

## RESULTS

### Antibody removal by 2ME/SDS

A method based on strong oxidant (beta mercaptoethanol; 2ME) and a detergent (sodium dodecyl phosphate; SDS) was previously published (20).

We applied this method to sequential staining and stripping over ten cycles with nine different antibodies representing membrane, cytoplasmic and nuclear proteins and quantified the staining results. Variations for each subsequent staining were comprised within 10% above or below the initial result at time 0 (Fig. 1 and Supplemental Fig. 3-A). We confirmed the reduced intensity for Ki-67 staining with 2ME/SDS incubation, however the change was about 5% of the initial value and did not changed after the first step (Supplemental Fig. 3-B). CD44 was similarly mildly affected by this stripping method.

**Fig. 1.**
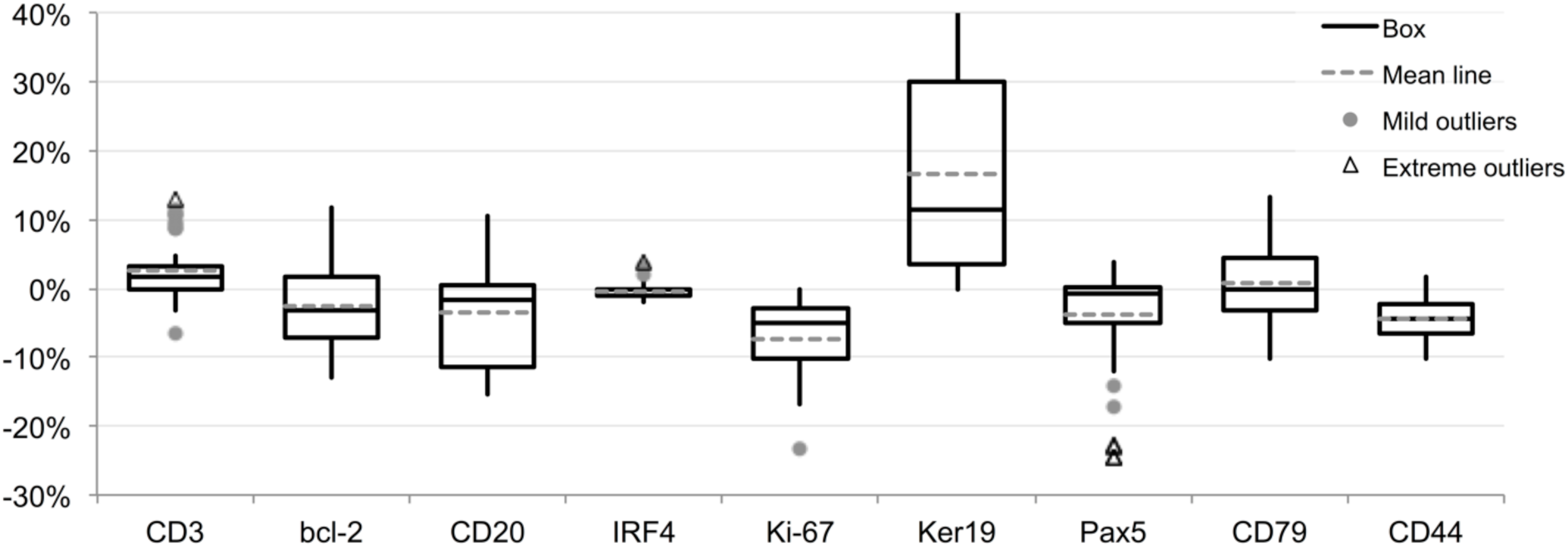
Variability for nine markers over ten staining and stripping 2ME/SDS cycles. One single section for every three markers was stained (t0) and sequentially stripped and re-stained for the same markers 10 times. Variation in staining intensity is expressed as fraction of the 256 8bit channels over the initial staining intensity. Primary Ab incubation time: 1 hr, 2nd Ab: 30 min, single indirect IF. Data are from four independent 10-cycle experiments.

An enhancing effect of the 2ME/SDS treatment, previously described (20) e.g. on keratins, was not apparent or reduced when the ARx (30) initial treatment was applied. ARx maximizes the immunoreactivity and overcomes the requirement for a tailored pH for AR (ibid). Most of the nine antigens are expressed homogeneously on the cell, therefore we digitally threshold the images and measured the amount of pixel, as a measure of the effect of the staining variations on the detection of the target. There was little variation in the amount of positive cells detected over the cycles (Supplemental Fig. 3-C).

### Antibody removal by chaotropic salts

The 2ME/SDS stripping method uses hazardous chemicals with a strong odour, therefore we investigated alternative methods.

Initial experiments of antibody removal with antichaotropic (saturated ammonium sulphate) and chaotropic compounds (GnHCl) showed that saturated ammonium sulphate was ineffective (not shown) but GnHCL at 5.3M and 6M efficiently removed primary and secondary antibodies from sections (Fig. 2). The process was more efficient at pH 4 (33) (not shown). Differently from 2ME/SDS, very dense spatially arranged proteins (e.g. pentameric cytoplasmic IgM, but not dimeric IgA or IgG) could not be completely stripped by GnHCl of bound primary and secondary antibodies (Fig. 2).This did not happened with 2ME/SDS.

**Fig. 2.**
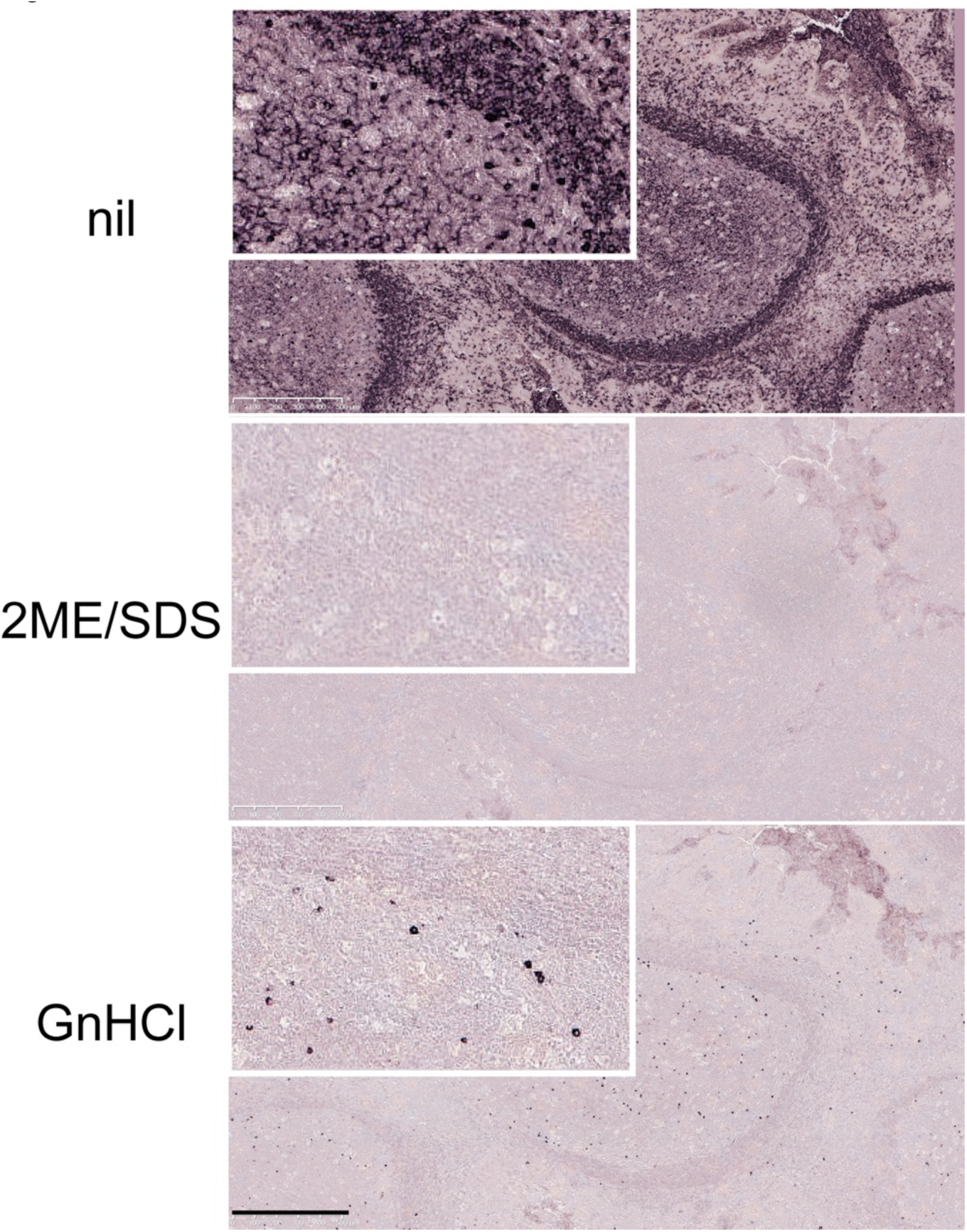
Comparison of antibody removal efficiency of 2ME/SDS and GnHCl Tonsil sections were incubated with a FITC-conjugated anti human IgM, stripped with either 2ME/SDS or GnHCl, stained with a FITC-conjugated anti-human IgM rabbit Ab, counterstained with an AP-conjugated anti-FITC antibody and developed in NBT-BCIP. 2ME/SDS stripping (middle) leaves no stainable primary antibody. Stripping by GnHCl (bottom) leaves IgM+ plasma cells. Identical results are obtained with an anti-Rabbit AP-conjugated (not shown). High magnification in the insets. Scale bar: 500 μm.

The treatment with GnHCl alone reduced the antigen availability (33): this negative effect could be fully reversed by transferring the sections to a 6M urea solution (6MU), followed by a 3M urea solutions (33), both containing sucrose as a protein refolding agent (ibid) and/or by a brief 1 min AR step (not shown). The combination of both 6M and 3M urea followed by a short AR treatment in 10 mM EDTA in Tris-buffer pH 8 was found the most effective.

Variations for each subsequent staining over ten stripping cycles were comprised between -5 % and +30% of the initial result at time 0 (Fig. 3). The enhanced staining obtained with some antigens after one or more stripping cycles is due to the short AR step, since the same antigens showed a reduced detection (+5% to -15% variations over five cycles) when the AR step was not included (Supplemental Fig. 4). However, longer AR after a GnHCL-6MU treatment may mildly decrease some antigens (e.g. CD20, CD44) (not shown). A broader variation in the amount of cell target detected was observed (Supplemental Fig. 3-C) compared with the data obtained with 2ME/SDS.

**Fig. 3.**
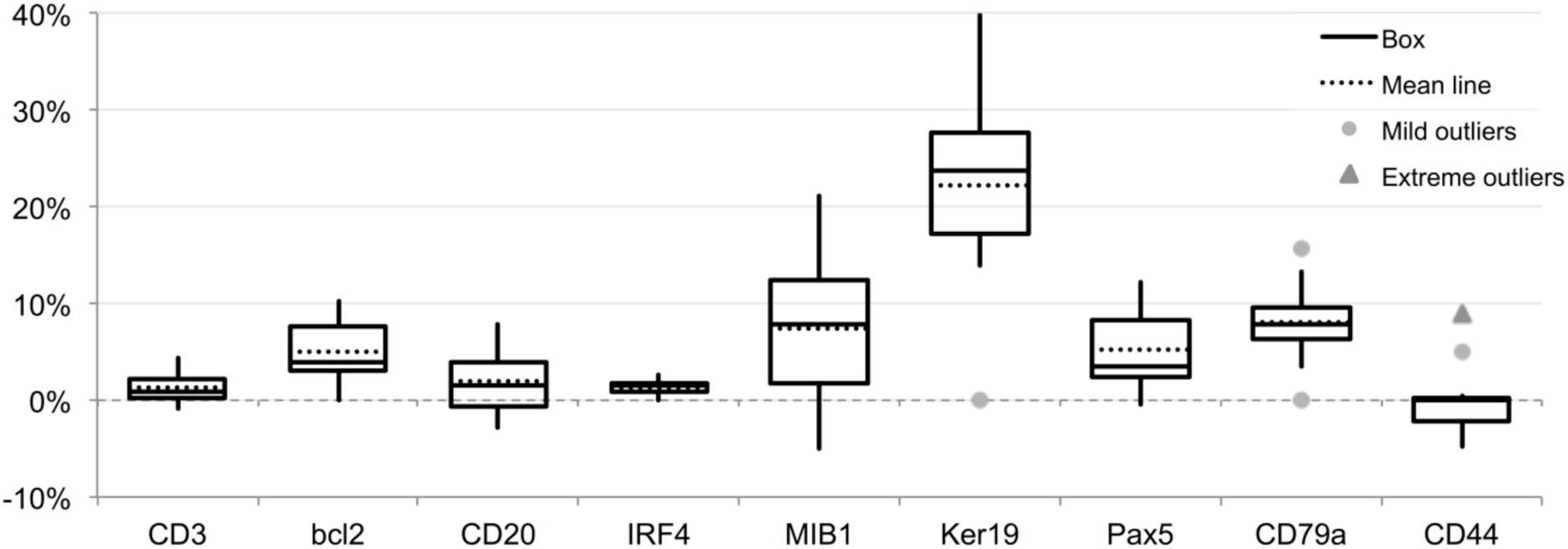
Variability for nine markers over ten staining and stripping GnHCl, 6Murea cycles. One single section for every three markers was stained (t0) and sequentially stripped and re-stained for the same markers 10 times. Variation in staining intensity is expressed as fraction of the 256 8bit channels over the initial staining intensity. Primary Ab incubation time: 1 hr, 2nd Ab: 30 min, single indirect IF.

### Antibody removal by Sodium Borohydride

Sodium Borohydride (NaBH_4_) has proteolytic properties (41), acting on peptide bonds mildly and selectively (42, 43), particularly for serine, threonine and asparagine, but not others. NaBH_4_ can cleave peptide linkages and reduce disulphide bonds (43). In addition, it suppresses fluorochrome fluorescence (44). Thus it is an attractive chemical to remove bound antibodies. We tested NaBH_4_ 15 mM on bound antibodies of different species and mouse isotypes and found it most effective on rabbit and goat Ig, less on mouse IgG1 and IgG2a (Fig. 4). NaBH_4_ at lower molarity was much less effective (not shown). The addition of 0.5% SDS improved the removal of mouse IgG2a and partially of rabbit and goat Ig, at the cost of tissue damage with repeated applications (not shown).

**Fig. 4.**
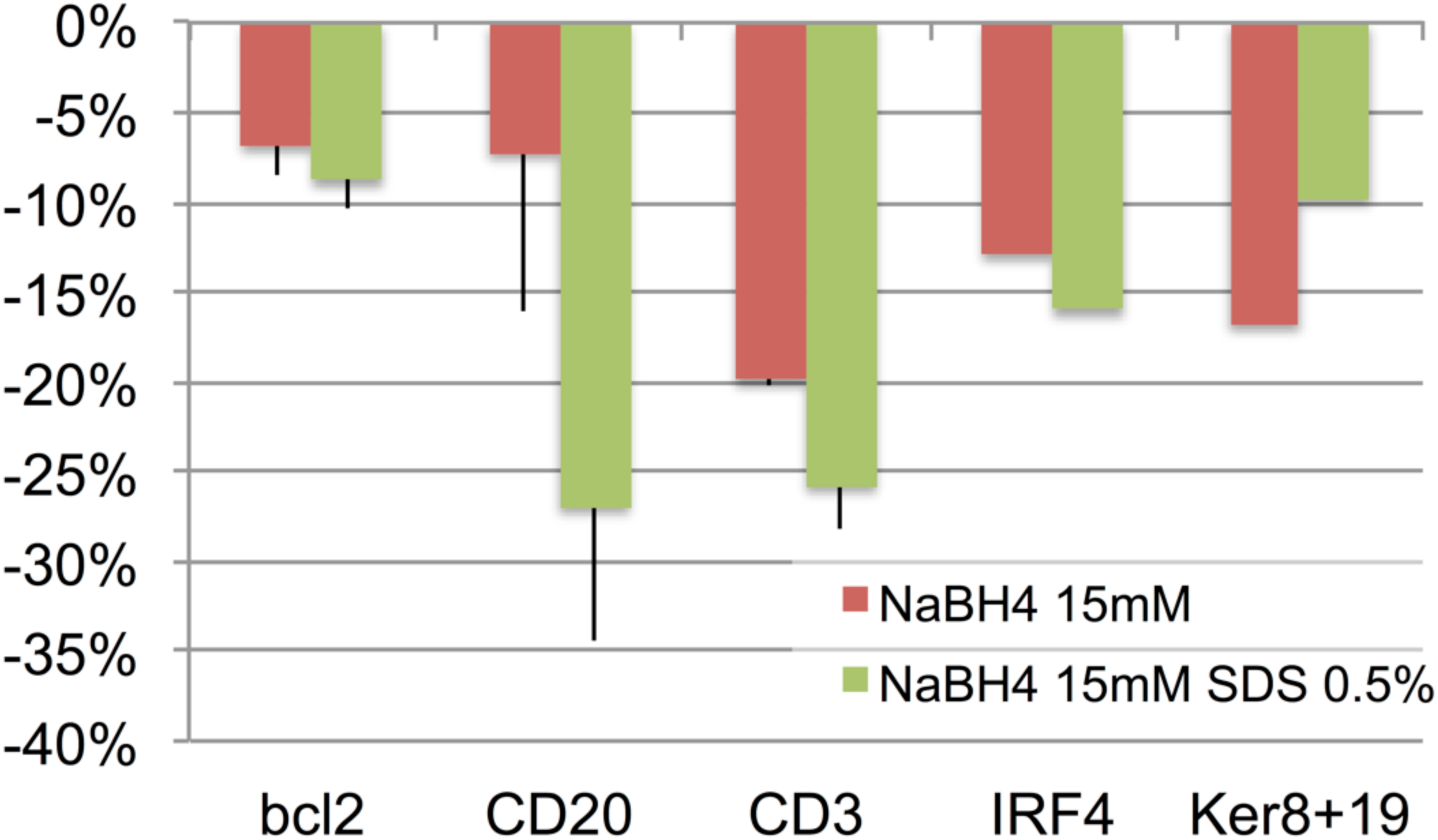
Effect of NaBH_4_ 15mM alone or in combination with SDS on bound primary antibodies. Stripping is enhanced by SDS and the effect is proportional to the starting abundance of the target. Bcl-2: mouse IgG1; CD20: mouse IgG2a; CD3: rabbit Ig; IRF4: goat Ig; Ker8+19: pooled rabbit Ig anti keratin 8 and 19.

The application of 1 mg/ml (26mM) NaBH_4_ in buffer at RT for longer than 20 min results in loss of the tissue (Supplemental Fig. 5-A, -B). However, NaBH_4_ treatment for 10 min at a concentration between 0.6 and 15 mM do not affect tissue antigens, even after repeated applications (Fig. 5). Because of its similarities with 2ME in reducing disulphide bonds, we tested NaBH_4_ as a 2ME substitute in the ARx method (30), and found it ineffective (not shown).

**Fig. 5.**
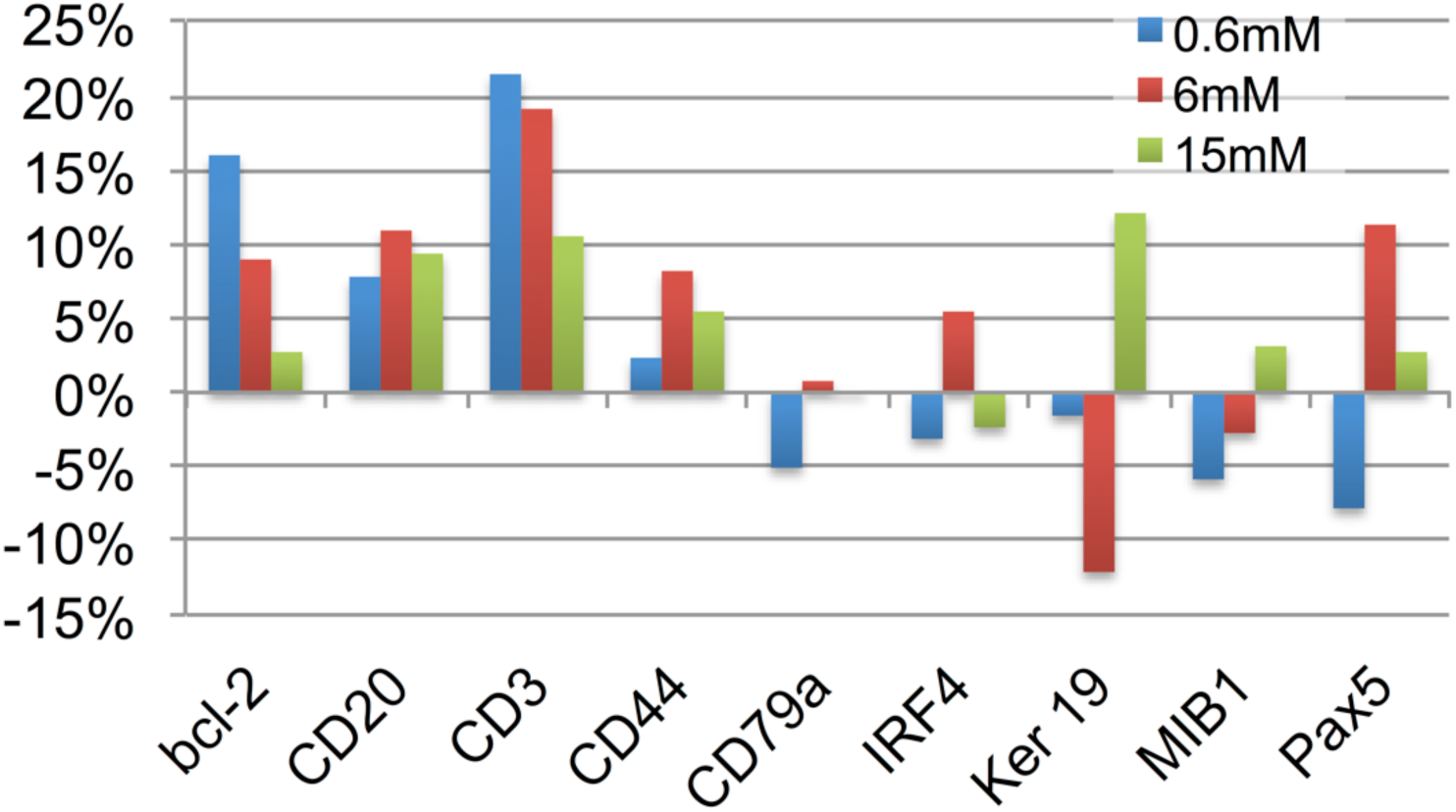
Effect of NaBH_4_ 0.6, 6 and 15mM on antigens. Sections have been treated with 10 cycles of 10 min NaBH_4_ pH9, followed by pH 4 buffer. Channel intensity variation over 256 channels is shown.

### Effect of antibody removal on target detection and AF

As shown in Fig. 1, 3 and Supplemental Figs. 3 and 4, the variation in staining over repeated cycles of staining and stripping is low. As published before (20), a decrease in staining affects only some antigens and here we show that it is limited to the very first cycles (Supplemental Fig. 3-B).

Moreover, by alternating positive antibody and negative control staining over ten cycles, the controls remain below the lower detection limit of a positive stain across the cycles (Supplemental Fig. 3-A)

To quantitatively assess the extent of antibody removal by 2ME/SD and GnHCl we first addressed how the two stripping methods affect the fluorochromes, in order to distinguish between antibody removal and fluorochrome quenching. Stained sections were extensively crosslinked with formalin and subject to 2ME/SD or GnHCl stripping; differently from NaBH_4_, no effect on fluorescence was observed after both stripping method (not shown). Formalin fixation did not affected fluorescence (not shown). Based on this result, any change in staining must be due to antibody removal.

We then stained in double indirect IF the sections for abundant proteins with repetitive motifs (i.e. keratins and vimentin), measured the staining intensity, stripped, re-stained the stripped sections with negative control Abs and fluorochrome-conjugated antibodies (Fig. 6 and Supplemental Fig. 6). Not only the antibody removal, both primary and secondary, was almost total, but re-staining with the same powerful double indirect IF failed to detect a significant amount of leftovers.

**Fig. 6.**
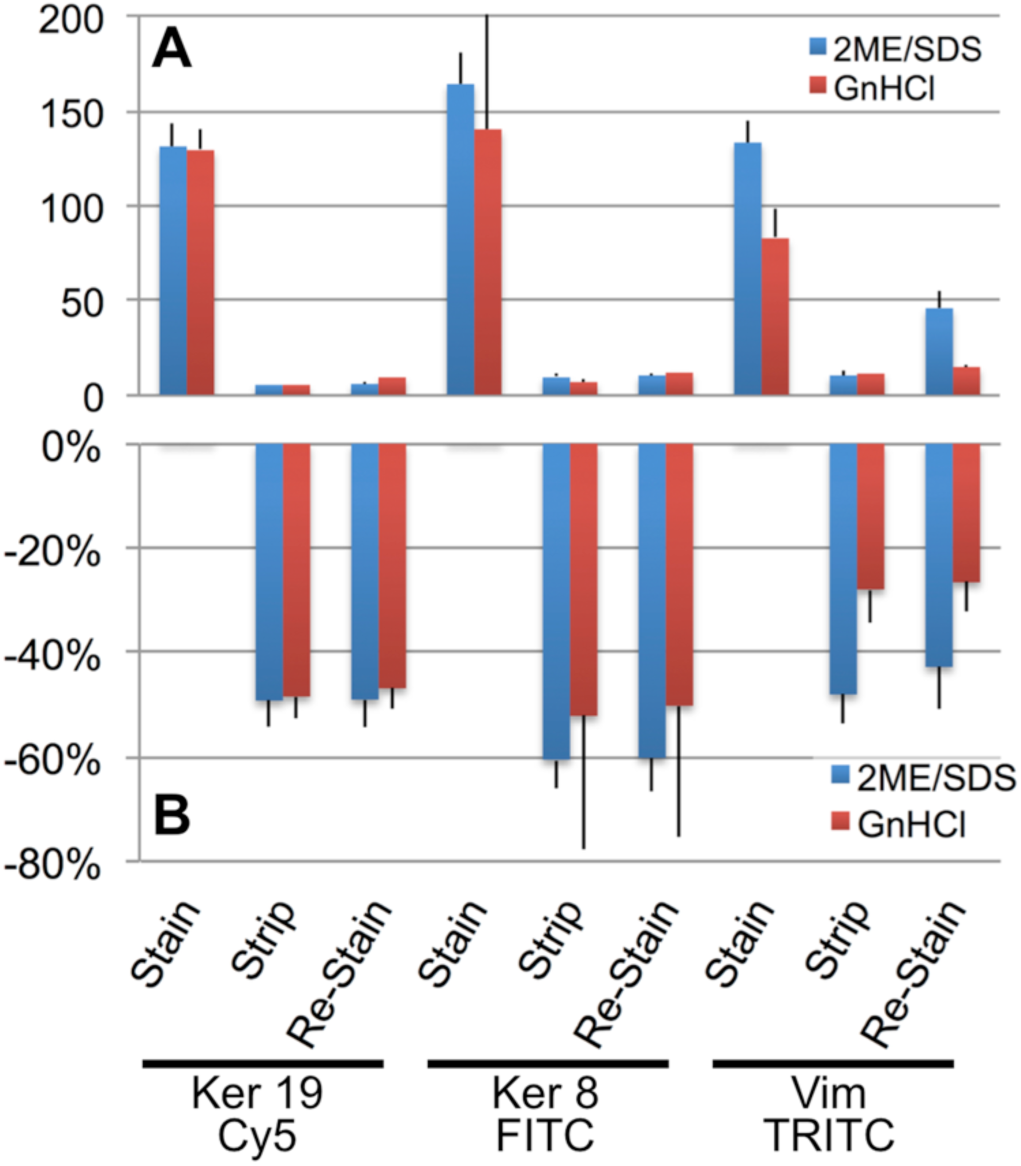
Effect of stripping and re-staining on abundant antigens. Sections in duplicate were stained in double indirect IF for the indicated abundant antigens (intermediate filaments). The channel value (A) or the % changes from the t0 stain over 256 channels (B) ± SD are plotted before (Stain), after stripping (Strip) and after re-staining with species- and isotype-matched non-immune control Abs (Re-Stain). Primary Ab incubation time: O/N, 2nd Ab: 30 min, double indirect IF.

We succeeded in sequentially staining and stripping in excess of 20 cycles (i.e. detection of 63 antigens) a single FFPE section, using the 2ME/SD method.

AF obtained before and after AR was qualitatively different, as measured by pixel-by-pixel comparison (Supplemental Fig. 2-E).

For most tissues, where AF is scarce, there was little variation over several staining and stripping cycles; however, AF-rich tissues such as kidney displayed a bimodal loss of AF over the cycles (Fig. 7). Pixel-by-pixel comparison at time 0 and after the last stripping showed an heterogeneous loss of AF, not seen in an immune infiltrate in the same section (Supplemental Fig. 2-F), suggesting selective inactivation or extraction of highly fluorescent tissue components.

**Fig. 7.**
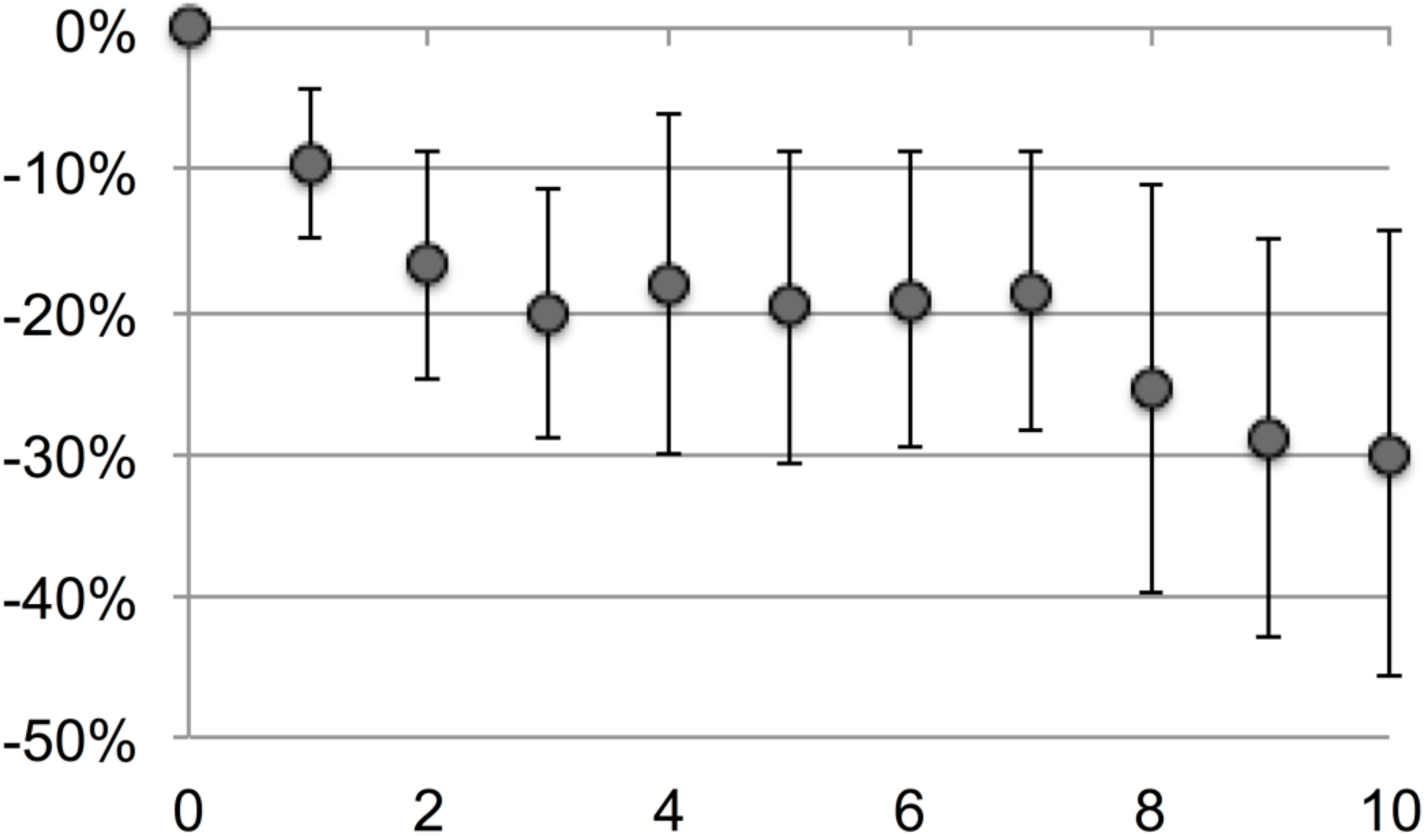
Cycle-dependent decrease of kidney tissue autofluorescence. Variation over time of the kidney tubules AF (em 520nm), expressed as % channel changes from time 0, over 10 staining and 2ME/SDS stripping cycles. Eleven different areas from three samples have been imaged and the average intensity change over the starting AF is expressed as % change over 256 channels ± SD

### Autofluorescence (AF) chemical subtraction

Exposure of highly autofluorescent tissues to NaBH_4_ 15 mM in 95% EtOH for 10 min reduced the tissue AF in the three channels by 6-16% below the control values (Supplemental Fig 5). The reduction was specifically due to NaBH_4_, since the vehicle, NaOH 0.05 N, produced a modest increase in AF. The treatment of dewaxed sections, before the AR step, for 10 min with 15 mM NaBH_4_ caused a 19-24% increase of AF and a significant reduction of antigenicity (not shown) and, if contacted for 30 min or more, partial or total loss of the tissue.

In multiple experiments on tonsil and kidney tissue, no significant AF reduction was obtained (not shown) as previously published (45).

## DISCUSSION

A dewaxed, antigen retrieved FFPE section is essentially identical to a Western Blot sheet (46), which can be stained and stripped multiple times of previously deposited antibodies. In a Western Blot the proteins are linearized and separated by molecular weight. The FFPE tissue, however, contains individual proteins crosslinked in situ with unknown bystanders whose epitopes are rescued from the processing-associated masking and re-exposed. As for the proteins immobilized on a membrane, tissue epitopes, once re-exposed, need to maintain the minimum amount of water in order to preserve the shape in an immuno-recognizable fashion (28). This essential requirement has been totally overlooked in routine IHC or IF, because there has never been the need to re-use a stained section, until it became a prominent issue for reproducibility when performing multiple sequential staining rounds. By moving the concept of a tissue section closer to a blot membrane and by examining the effect of every component on the antigens at each step of the mplx process, we have achieved optimal reproducibility and a low coefficient of variation upon repeated staining.

### Diverse molecular mechanism of antibody removal yield an identical outcome

The two methods shown here for multiplexing work with two quite different mechanisms. The 2ME/SDS method is chemically altering the structure of the primary and secondary antibodies. Disulphide bonds are in a thermodynamic equilibrium for each given type of antibody (47): the strong reducing agent, 2ME tilts the balance toward an unbound form. The detergent SDS favors the dissociation of the Ig heavy chains (48) and the combined effect overcomes the bond between epitope and paratope, even for high-energy affinity (20). The heavily crosslinked tissue is resistant to the combined action of 2ME and SDS, at least for repeated exposure but at a temperature below the one required for tissue solubilization (30).

The decrease of tissue AF with stripping cycles is in contrast with the little variation shown in antigen detection. Data for the smallest protein tested, bcl-2 weighing 26 kDa, point to a stable presence over repeated cycles for proteins of that molecular weight or larger, crosslinked in the tissue. Either repeated application of 2ME and SDS are quenching the AF, or small soluble autofluorescent compounds, not firmly crosslinked in the tissue, may be progressively removed from the section. A similar decrease of autofluorescence with cycle repetition was observed with a fluorochrome-quenching technique (49); differently from this method, we did not observed cell loss.

The method based on GnHCL produces a rapid conformational change of the epitope and of the paratope (50), in part altering the water structure which holds the epitope in shape. The detachment of the bound antibody is the effect sought, but the amount of the available epitopes is significantly diminished, probably because of incorrect re-folding of epitopes after GnHCl (33). In order to correctly re-fold any available epitopes exposed after AR, the section is transferred to high molar urea (33), then to a diluted urea solution, in the presence of a folding enhancer, sucrose (51). Epitope-refolding is further enhanced with a brief exposure to high temperature, analogously to AR.

The interaction of GnHCl with the protein-associated water molecules in the first phase of denaturation (50) may account for the inability to remove antibodies bound in high amount in packed molecular structures, possibly resulting in precipitation of the antibody at the antigen site and insolubility, because of the removal of critical water molecules.

The use of a chaotropic agent to reversibly modify the epitope conformation in a controlled fashion, in order to detach a bound antibody, further emphasizes the metastable nature of re-exposed epitopes in FFPE material, as previously published (28). The efforts aimed at preserving the epitope conformation during each and every step of the mplx process, including antibody incubation and coverslipping, represent the other novelty of the present method: reproducibility over multiple staining cycles has been obtained by using buffers and chemicals tested on purpose by quantitative IF for epitope stability. Differently from other mplx methods (26), removal of antibodies with 2ME/SDS do not requires additional treatments such as AR between cycles, leaving tissue antigens unmodified for the whole sequence. Because of that, no staining prioritization is necessary.

### Comparison of 2ME/SDS vs GnHCl

Despite the identical outcome of the two protocols, there are specific characteristics for each one, which suggest a preferential use.

GnHCl is a non-toxic, safe method which works best where a short cycling sequence is required and sparsely represented proteins are involved. The enhancing effect of the short AR step may be exploited to prioritize the staining sequence. The inability to entirely remove dense antibody deposits such as IgM in plasma cells is compensated by the poor immune reactivity of the aggregated leftovers which are eventually left (Fig. 2).

The 2ME/SDS method has a thorough antibody removal ability and, combined with disaccharide protection, minimal or no effect on the antigens over extensive staining and stripping cycles. However, it depends on an optimal antigen retrieval from the very beginning, because no additional AR steps are involved. It requires a waterbath with controlled temperature and confinement of a chemical with a strong odor.

The extent of primary and secondary antibody removal was tested in multiple ways, quantitative and qualitative, exemplified in Fig. 2, Fig. 6, Supplemental Fig. 3-A and 6. First we excluded a bleaching effect of both 2ME/SD and GnHCl on the fluorochromes used. Next we acquired the tissue images post-stripping with the very same fluorescence setting. Finally, we re-stained stripped sections with the same protocol used to get the initial stain, with the substitution of a negative, isotype matched irrelevant antibody as the first step (Fig. 6, Supplemental Fig. 3-A and Fig 6) and imaged again. To test for the consistency of stripping, sections were consecutively stained with a positive and a negative antibody every other cycle (Supplemental Fig. 3-A for 2ME/SDS and not shown for GnHCl). To test for stripping completeness in IHC (Fig. 2), we used a small molecule (FITC) as a robust hapten, resistant to modifications (28) and a sensitive IHC development. The aggregate results show that residual antibodies after stripping, if ever present, are below detectability, within the staining method employed.

During this investigation, we came across multiple interesting characteristics of sodium borohydride, a compound used in biochemistry, fluorescent microscopy and tissue fixation. The ability to quench the fluorochrome emission and the proteolytic activity suggested its use for multiplexing, although we found that in several previous reference papers the pH of the diluent used shortened the half-life and therefore its action to mere seconds. Differently from proteases, occasionally used for multiplexing (8, 23) and known to affect tissue antigens, NaBH_4_ by itself proved to possess some ability to remove antibodies from tissues without damage (Fig. 4 and 5). However the addition of SDS causes substantial damage to the tissue with treatment repetitions and NaBH_4_ was not further pursued.

### Antibody removal-based multiplexing is a powerful technique of widespread use

There are several advantages for the antibody stripping method we describe.

All reagents and instruments are common, commercially available, low cost, compared with other techniques employing directly conjugated primary antibodies, amplification kits, complex software and spectral microscopes.

One single section of routine FFPE material is required, opening a wealth of clinical specimens for study. Since all the stains are eventually combined in one multiplexed stack of images, the system can be optimized by selecting antibodies which differ by species and isotypes, filling each staining round with three stains.

Steric hindrance caused by interacting or heterodimerizing proteins can be circumvented by spacing the stainings in two separate staining cycles.

In addition, mistakes in staining can be mended by re-staining for the same antibody.

Despite the fact that the filter setting we used allows the detection of seven fluorochromes and up without crosstalk (DAPI / BV421, FITC, TRITC, Cy5, Pacific Orange / BV480, BV605, PerCp-Cy5.5, BV711; see <u>https://goo.gl/scE3sd</u>), the limit is the simultaneous detection in indirect IF of two antibodies of the same kind with two separate fluorochromes, an area we are investigating.

During this investigation we found convenient to allocate antigens in the various channels according to abundance (lowest to highest) in the following order: TRITC, Cy5, FITC. This because of the dye brightness with our filter set (Rhodamine RedX: 1.55 x 10^-6^ mW; Alexa 488: 2.89 x 10^-7^ mW; Alexa 647: 3.96 x 10^-7^ mW), the differential sensitivity of the sensor to light (maximal between 520 and 640 nm), the absence of AF in the Cy5 channel, the digital removal of AF.

The combination of *i)* a careful choice of validated antibodies used at a concentration not exceeding 1μg/ml, *ii)* an enhancements of the sensitivity (superior AR, double indirect IF), *iii)* the choice of buffers and mounting media, *iv)* the acquisition with a sensitive, linear detector, *v)* the removal of the noise caused by AF, all concur to deliver a superior signal over multiple staining and stripping cycles.

A limitation of some popular mplx methods consists in the size limit of the area that can be digitally acquired, either because of how the software has been designed or because of long acquisition time or the image format. By acquiring whole slide images in a format that can be read by many platforms, the areas of interest need not to be known upfront and additional regions of interest can be investigated later. Care should be taken to increase the computing RAM to at least 32 Gb and to limit each single whole slide image below 6 Gb.

### Multiplexing as a discovery tool

Mplx IF as we describe here is a powerful discovery tool. By applying as little as a dozen markers on the very same tissue section, we could dissect the phenotype of rare and scattered immune cells such as interfollicular dendritic cells or germinal center follicular helper T cells, highlighting the power of multiplexing to investigate polymorphous reactive immunopathology. As shown in Fig. 8, S-100+ cells of dendritic appearance can be simultaneously assessed for activation, proliferation, Class II MHC and transcription factors expression, monocyte markers, and geographic localization in the tissue.

**Fig. 8.**
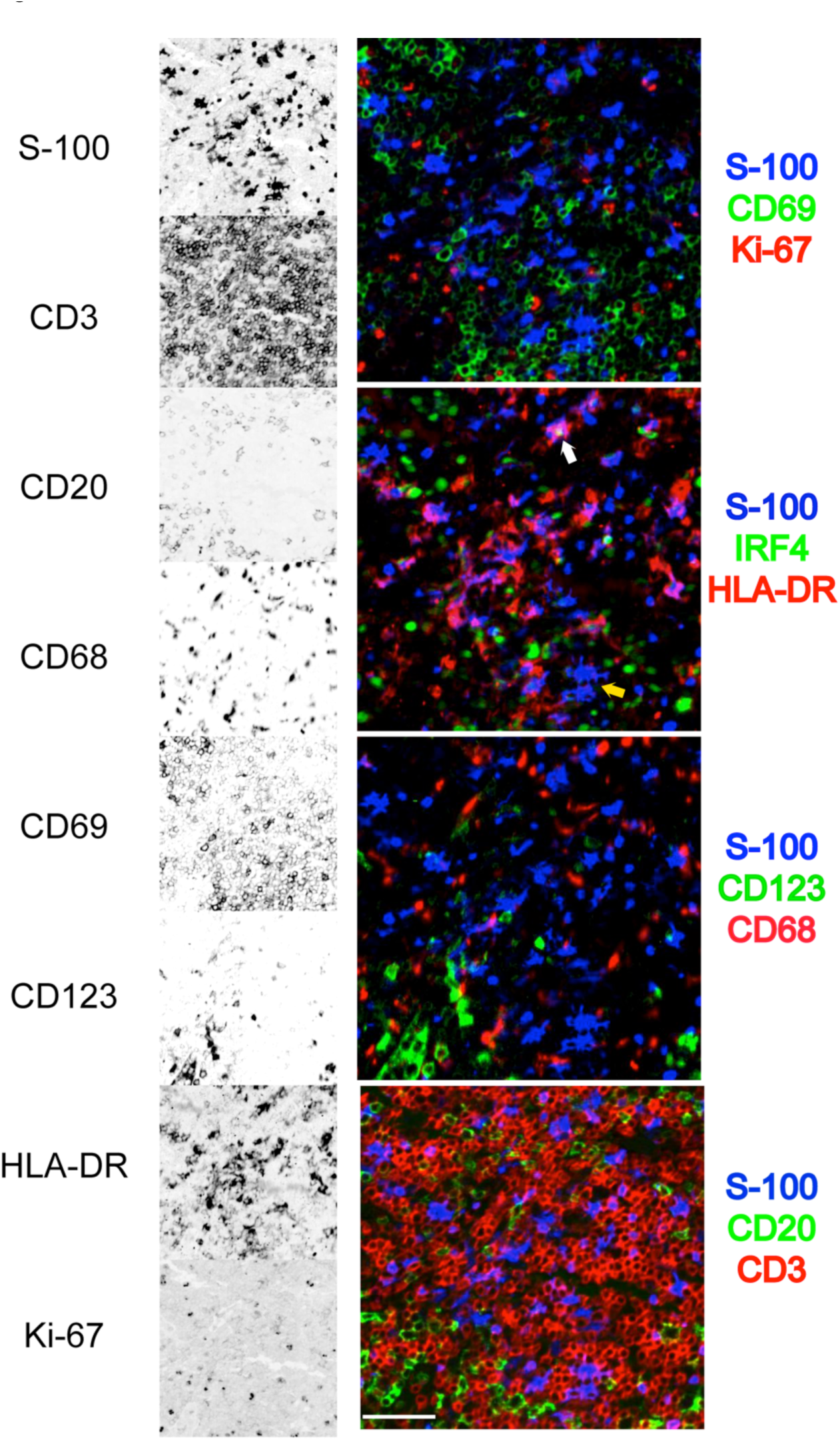
Detail of a multiplexed interfollicular tonsil area. Eight stains out of a 32 antibody multiplex are selected from a tonsil interfollicular area and shown as single stains (inverted grayscale) on the left, or combined into four three color RGB composites. The area contains numerous S-100+ dendritic cells, HLA-DR+ with exceptions (yellow arrow), occasionally IRF4+ (white arrow), negative for Ki-67, CD69, CD68 and CD123. The dendritic cells are located in a CD3+ T-cell area, without contact with CD20+ B cells. GnHCl stripping method. Scale bar = 100 μm

By multiplex staining, we re-assessed the expression of PD-1/ CD274 on a subset of germinal center proliferating B cells expressing BCL6 (Fig. 9), a previously described subset (52). Interestingly, we were able to confirm the B cell expression by using the UMAB197 IgG2a but not the NAT105 IgG1 antibody, despite the full co-localization on follicular helper T cells (not shown).

**Fig. 9.**
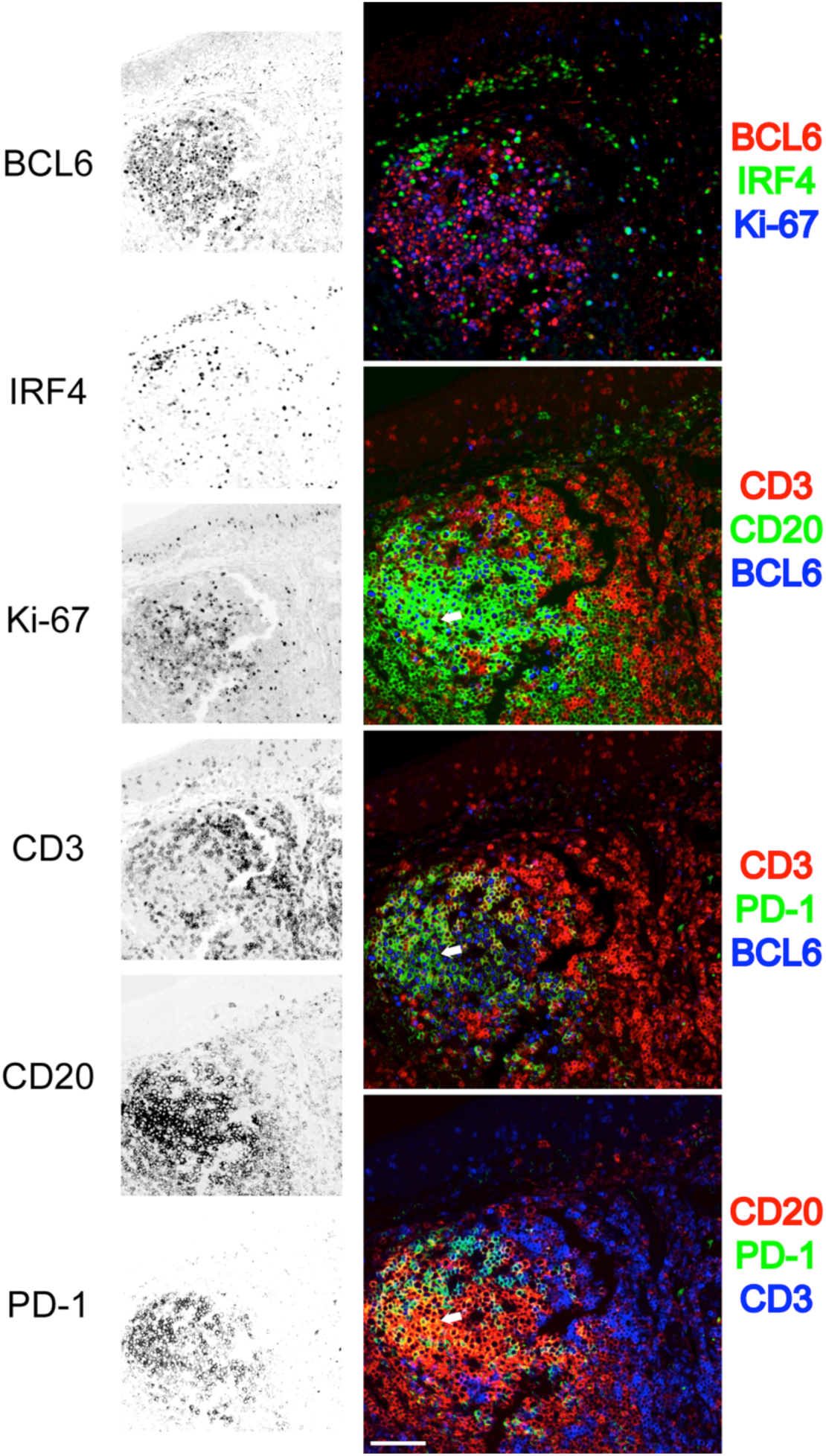
Detail of a multiplexed tonsil follicle. Six stains out of a 32 antibody multiplex are selected from a tonsil follicle and shown as single stains (inverted grayscale) on the left, or combined into four three color RGB composites. Proliferating (Ki-67+) germinal center BCL-6+ CD20+ B cells are mutually exclusive stained for IRF4. PD-1 stains CD3+ T cells in the germinal center as well as rare PD-1+, BCL6+, CD20+ centroblasts (white arrow). GnHCl stripping method. Scale bar = 100 μm

The comparison in cell number detection shown here (Supplemental Fig. 3-C) between traditional single color IHC and multiplex IF opens the use of multiplexing for clinical investigation and use.

## AUTHOR’S CONTRIBUTIONS

Giorgio Cattoretti, Carla Rossana Scalia and Maddalena Maria Bolognesi equally designed the experiments.

Carla Rossana Scalia performed immunofluorescent tests, acquired the digital preparations. Maddalena Maria Bolognesi and Marco Manzoni devised the image analysis algorithms. Francesca Maria Bosisio provided essential reagents.

Mario Faretta provided essential knowledge on fluorochromes characteristics, filter settings and image analysis tools.

Francesca Maria Bosisio and Stefano Zannella performed visual and digital image analysis. Giorgio Cattoretti and Maddalena Maria Bolognesi wrote the manuscript.

All authors have read and approved the final manuscript.

## ACKNOWLEDGEMENTS

We wish to thank Dr. Franco Ferrario for continuous support, Emanuele Martella, Marco Cicuttin (Nikon, Italia) and Giulio Simonutti (Hamamatsu Italia) for expert advice, Linde DeSmedt (HistoGeneX NV, Antwerpen, Belgium) for suggestions, Riccardo Tagliabue for expert technical help.

The Hamamatsu S60 digital scanner was obtained as part of a clinical research project BEL114054 (HGS1006-C1121) of the University of Milano-Bicocca and GlaxoSmithKline, on which project Carla Rossana Scalia and Maddalena Maria Bolognesi are also supported.

## SUPPLEMENTAL TABLE LEGENDS

Supplemental Table 1: Primary antibodies

Supplemental Table 2: Secondary antibodies

Supplemental Table 3: Staining variability

The mean channel value ± SD is shown for the number of replicas in the column “n”. Data are expressed also as % variation over 256 channels. The wavelength in which the variation is measured is also shown

## SUPPLEMENTAL FIGURE LEGENDS

Supplemental Fig. 1

A-Effect of incubation time on staining

The variation in intensity, measured as channel variation, after one hour (100%) and overnight (O/N) incubation is plotted.

B-Effect of double indirect vs single indirect staining of intensity.

The variation in intensity, measured as channel variation, after one (100%) or two rounds of indirect immunofluorescent staining are plotted.

C-Effect of mounting media on antigenicity.

Buffered 60% Glycerol as mounting medium has minimal or no effect on antigen preservation, compared to a polyvinyl alcohol or a Glycerol-Gelatin one. Antigen-retrieved sections were mounted with the media shown, immunostained and quantified. Values are % channel variations over the baseline (100%).

Supplemental Fig. 2

A-Tissue autofluorescence in four tissues.

Tissue autofluorescence was obtained by multiple exposure time in four tissues (placenta, kidney, skin and squamous epithelium) in each of four excitation/emission combinations. Mean fluorescence intensity (Y) is plotted against exposure time (milliseconds, X). Note the linear response.

B-A portion of a kidney biopsy stained for cytokeratin 8 was acquired with the AF (top) and with the FITC (middle, KER8+AF) specific filter sets. The AF image, corrected for the exposure factor, was subtracted from the cytokeratin image. The result is shown at the bottom, showing exclusively the specific staining (KER8). Note the absence of nuclear DAPI staining in the top image. Scale bar = 500 μm.

C and D-Pixel-by-pixel comparison of AF values in placenta and kidney.

Comparison of the intensity values pixel by pixel are shown for the AF channel (420exc/530em) versus the FITC and the TRITC channels in placenta (left) and kidney (right). FITC vs TRITC comparison is also shown. Note in kidney the spread of the values for AF vs TRITC and for FITC vs TRITC.

E-Changes in autofluorescence before AR, after AR and after stripping (5 cycles). Autofluorescence images were obtained from placenta sections, illuminating with the 480 ± 17nm filter and collecting with the triband dichroic and the 520 ±28nm filter. WSI were acquired before AR, after AR and after five cycles-equivalent (2,5 hr) of 2ME/SDS stripping. The WSI were all registered and a 30,000 pixel selection placed in an excel file. Bivariate plot for PreARPostAR (red dots) and PostAR-PostStrip (black circles) are shown. Note the multiple pixel populations with divergent AF before and after AR, while before and after stripping the vast majority of pixels line up along the middle of the bivariate graph.

F-Pixel-by-pixel comparison of AF values over 10 stripping cycles.

Comparison of AF values of kidney tubules (left) and an immune infiltrate (right) at time 0 (abscissa) and after 10 cycles (ordinate) shows an overall reduced AF values in the immune cells and a greater variation of the pre- and post-values in kidney tubules.

Supplementary Fig. 3

A-Channel intensity for six markers and six negative controls over ten staining and stripping 2ME/SDS.

One single section for every three markers was stained (t0), sequentially stripped with the 2ME/SDS method and re-stained alternatively with a negative control antibody or for the same markers 10 times. Staining intensity variation is expressed as a box plot. Note the separation of the negative controls from the positive stains. CD3 and Pax5 in FITC, bcl-2 and CD79a in TRITC, CD20 and CD44 in Cy5. Primary Ab incubation time: 1 hr, 2nd Ab: 30 min, single indirect IF.

B-Channel variations for individual antibody pools over stripping.

Percent variations from the t0 channel position over 256 channels for each three-antibody pool at each stripping cycle with the buffers listed on top of each column.

C-Variation of the positive pixel area for ten stripping cycles.

The positive pixel area, expressed as percentage of the area analyzed, over ten staining and stripping cycles is depicted for GnHCl (blue boxes), 2ME/SD (red boxes) and DAB-stained serial sections (black boxes). Note the overlap of the results over the three methods for most, but not all antigens.

Supplemental Fig. 4

Variability for nine markers over ten staining and stripping GnHCl, 6Murea, no AR cycles. One single section for every three markers was stained (t0) and sequentially stripped and restained for the same markers 10 times. Variation in staining intensity is expressed as fraction of the 256 8bit channels over the initial staining intensity. Primary Ab incubation time: 1 hr, 2nd Ab: 30 min, single indirect IF. Note the decrease below the baseline value for all the markers tested.

Supplemental Fig. 5

A, B-Effect of 26mM NaBH_4_ in EtOH 95% and Tris Buffer 0.05M pH 9.

NaBH_4_ causes changes in AF when applied at 1 mg/ml (26mM) in 95% EtOH (A) or in buffer pH9 (B). The reduction at 30 min in buffer is a NaBH_4_-specific degradation of the section. Changes in AF in three filter combos are expressed as % over 256 channels.

C-Effect of 15mM NaBH_4_ in EtOH 95% and Tris Buffer 0.05M pH 9 on AF.

A NaBH_4_ specific effect is seen in EtOH at 10 min and disappears at 30 min. The reduction at 30 min in buffer is a NaBH_4_-specific degradation of the section.

NaBH_4_ is applied at 15mM in 95% EtOH or in buffer pH9. An equivalent amount of NaOH, the diluent, is applied as a comparison. The reduction at 30 min in buffer is a NaBH_4_-specific degradation of the section. Changes in AF in three filter combos are expressed as % over 256 channels.

Suppl. Fig 6

Residual anti-intermediate filaments (keratins 8 and 19, Vimentin) primary antibodies after GnHCl and 2ME/SDS stripping

Selected high-power fields are shown from a kidney section stained for rabbit anti Keratin 8, mouse IgG2a anti Keratin 19 and mouse IgG1 anti Vimentin, before (top, control) and after stripping. Note that the control images are unmodified, acquired with the exposure time selected. The stripped images instead, taken with the same setting, have been modified with the automatic increase in brightness and contrast of the ImageJ software (Maximum value <20, Brightness ~ -10)

